# Antibodies against SARS-CoV-2 control complement-induced inflammatory responses to SARS-CoV-2

**DOI:** 10.1101/2023.05.29.542735

**Authors:** Marta Bermejo-Jambrina, Lieve E.H. van der Donk, John L. van Hamme, Doris Wilflingseder, Godelieve de Bree, Maria Prins, Menno de Jong, Pythia Nieuwkerk, Marit J. van Gils, Neeltje A. Kootstra, Teunis B.H. Geijtenbeek

## Abstract

Dysregulated immune responses contribute to pathogenesis of COVID-19 leading to uncontrolled and exaggerated inflammation observed during severe COVID-19. However, it remains unclear how immunity to SARS-CoV-2 is induced and subsequently controlled. Notably, here we have uncovered an important role for complement in the induction of innate and adaptive immunity to SARS-CoV-2. Complement rapidly opsonized SARS-CoV-2 via the lectin pathway. Complement-opsonized SARS-CoV-2 efficiently interacted with dendritic cells (DCs), inducing type I IFN and pro-inflammatory cytokine responses, which were inhibited by antibodies against the complement receptors (CR)3 and CR4. These data suggest that complement is important in inducing immunity via DCs in the acute phase against SARS-CoV-2. Strikingly, serum from COVID-19 patients as well as monoclonal antibodies against SARS-CoV-2 attenuated innate and adaptive immunity induced by complement-opsonized SARS-CoV-2. Blocking the FcyRII, CD32, restored complement-induced immunity. These data strongly suggest that complement opsonization of SARS-CoV-2 is important for inducing innate and adaptive immunity to SARS-CoV-2. Subsequent induction of antibody responses is important to limit the immune responses and restore immune homeostasis. These data suggest that dysregulation in complement and FcyRII signalling might underlie mechanisms causing severe COVID-19.

## Introduction

Since severe acute respiratory syndrome coronavirus 2 (SARS-CoV-2) was first identified in Wuhan, China, in December of 2019 [1, 2] the virus has spread all over the world, causing a respiratory disease, termed coronavirus disease 2019 (COVID-19) [3]. To date COVID-19 pathogenesis is still unclear. Asymptomatic patients and patients with mild COVID-19 gain control of infection within a couple of days most likely via innate immune responses as effective adaptive immune responses are expected to be elicited after 2 weeks in naïve individuals [4, 5]. Failure of antiviral innate responses to control infection might lead to uncontrolled viral replication in the airways eliciting an inflammatory cascade observed in severe COVID-19 cases [6, 7]. Severe to fatal outcomes in COVID-19 patients have been attributed to the dysfunction of innate and adaptive immune response by SARS-CoV-2 [8]. This aberrant or uncontrolled innate and/or adaptive immune responses lead to delayed viral clearance, inflammation and tissue damage, affecting organs [8–10]. It remains however unclear how the interplay of innate immune responses with adaptive immunity controls infection and how homeostasis is achieved after infection to prevent aberrant systemic inflammatory responses observed in severe COVID-19 disease.

The complement system constitutes an important innate immune response and acts as a first line of defence against viruses and might have a critical role in COVID-19 pathogenesis [11–14]. Complement activation limits SARS-CoV-2 infection but uncontrolled activity can lead to aberrant inflammatory responses observed during severe COVID-19 [11, 15, 16]. SARS-CoV-2 infection can activate complement by direct interaction of spike proteins with the lectin pathway via mannose-binding lectin (MBL) [17–20]. Interestingly, SARS-CoV-2-specific antibodies binding to spike protein also activate complement by the classical pathway through C1q [21, 22]. Moreover, the alternative pathway is triggered by SARS-CoV-2 spike protein by binding to cell surface heparan sulfates [23, 24]. Severe COVID-19 patients have high circulating C5a in their blood as well as high levels of processed C3 [16, 25], suggesting that uncontrolled complement activation might be involved in severity of COVID-19 [11, 26]. These studies suggest that although complement system is vital in limiting SARS-CoV-2 infection, dysregulation or lack of control of complement activation leads to severe pathogenesis [14, 26–28]. Mechanisms underlying complement-induced immunity and subsequent return to homeostasis after complement activation remain unclear.

Activation of mucosal dendritic cells (DCs) is a crucial step in the induction of effective innate and adaptive immune responses against invading viruses [29]. Notably, SARS-CoV-2 infection does not lead to strong DC activation [30–32]. Exposure of DCs to SARS-CoV-2 does neither lead to infection nor production of type I IFN and cytokine responses [33]. Although infection of bystander cells with SARS-CoV-2 can lead to DC activation [30, 34, 35], it is unclear whether complement deposition on SARS-CoV-2 can induce DC activation.

Here we investigated the role of complement in induction of immunity and how the inflammatory responses are controlled to prevent aberrant inflammation. Complement-opsonized SARS-CoV-2-induced DC maturation and efficient type-I IFN responses via complement receptors CR3/CD11b and CR4/CD11c. Moreover, complement-opsonized SARS-CoV-2 induced pro-inflammatory cytokines as well as IL-1β by caspase-1 inflammasome activation. Notably, serum from COVID-19 patients or anti-SARS-CoV-2 antibodies abrogated complement-induced DC activation and subsequent type I IFN and cytokine responses via CD32 activation. These data strongly suggest that complement is important in induction of innate and adaptive immunity but that antibody responses either elicited after infection or vaccination suppress complement-induced immunity and restore homeostasis. These data strongly suggest that antibodies against SARS-CoV-2 might be important in switching off complement-induced immunity and could be used to treat patients suffering from severe COVID-19 [36–39].

## Materials and methods

### Ethics statement

This study was performed according to the Amsterdam University Medical Centers, location Academic Medical Center (AMC) and human material was obtained in accordance with the AMC Medical Ethics Review Committee (Institutional Review Committee) following the Medical Ethics Committee guidelines. This study, including the tissue harvesting procedures, was consent by all donors and conducted in accordance with the ethical principles set out in the declaration of Helsinki and was approved by the institutional review board of the Amsterdam University Medical Centers and the Ethics Advisory Body of the Sanquin Blood Supply Foundation (Amsterdam, Netherlands). All research was performed in accordance with appropriate guidelines and regulations.

### Patient consent

To enable comparison between complement and antibody response following infection, we include serum collected in the RECoVERED cohort [40]. In total 10 RECoVERED serum samples from participants who experienced mild (5x) or severe (5x) COVID-19. Written informed consent was obtained from each study participant. The study design was approved by the local ethics committee of the Amsterdam UMC (Medisch Ethische Toetsingscommissie [METC]; NL73759.018.20). All samples were handled anonymously.

### Reagents and antibodies

The following antibodies were used (all anti-human): CD86 (2331 (FUN-1), BD Pharmingen), CD80 (L307.4, BD Pharmingen), PE-conjugated mouse IgG1 CR3/CD11b (101208, Biolegend), LEAF purified CR3/CD11b mouse IgG1, LEAF purified CR4/CD11c mouse IgG1, CR3/CD11b (M1/70), CR4/CD11c (S-HCL-3), CD32 (FUN-2), DC-SIGN (FAB161F, R&D systems), CD16, CD32 CD64 (all BD Pharmingen) and, viability dye (Ghost DyeTM Violet 510, Tonbo Biosciences, San Diego, USA). For extracellular staining, cells were incubated in 0.5% PBS-BSA (Sigma-Aldrich) containing antibodies for 30 min at 4°C. Single cell measurements were performed on a Canto flow cytometer II (BD Biosciences) and data was analyzed using FlowJo V10.8.1 (Software by TreeStar). Neutralizing COVA1-18 and non-neutralizing COVA1-27 antibodies were isolated from participants in the “COVID-19 Specific Antibodies” (COSCA) study [41] and were generated by Karlijn van der Straten as described previously [41]. Pre-pandemic normal human serum (NHS) generated from a pool of 10 healthy individuals was stored at -80°C until use.

### Cell lines

The simian kidney cell line VeroE6 (ATCC® CRL-1586™) were cultured in CO_2_ independent medium (Gibco life Technologies, Gaithersburg, Md.) supplemented with 10% fetal calf serum (FCS), L-glutamine and penicillin/streptomycin (10µg/mL). Cultures were maintained at 37°C without CO_2_. The human embryonic kidney 293T/17cells (ATCC, CRL-11268) were maintained in Dulbecco’s modified Eagle’s medium (Gibco Life Technologies) containing 10%FCS, L-glutamine, and penicillin/streptomycin (10µg/mL)

### DC generation

Human CD14^+^ monocytes were isolated from the blood of healthy volunteer donors (Sanquin blood bank) and subsequently differentiated into monocyte-derived DCs. In short, the isolation from buffy coats was performed by density gradient centrifugation on Lymphoprep (Nycomed) and Percoll (Pharmacia). After Percoll separation, the isolated CD14^+^ monocytes were differentiated into monocyte-derived DCs within 5 days and cultured in RPMI 1640 medium (Gibco Life Technologies, Gaithersburg, Md.) containing 10% FCS, penicillin/streptomycin (10 μg/mL) and supplemented with the cytokines IL-4 (500 U/mL) and GMCSF (800 U/mL) (both Gibco) [42]. After 4 days of differentiation, DCs were seeded at 1 × 10^6^/mL in a 96-well plate (Greiner), and after 2 days of recovery, DCs were stimulated or infected as described below. This study was performed in accordance with the ethical principles set out in the Declaration of Helsinki and was approved by the institutional review board of the Amsterdam University Medical Centers, location AMC Medical Ethics Committee and the Ethics Advisory Body of Sanquin Blood Supply Foundation (Amsterdam, Netherlands)

### SARS-CoV-2 pseudovirus production

For production of single-round infection viruses, human embryonic kidney 293T/17 cells (ATCC, CRL-11268) were co-transfected with an adjusted HIV backbone plasmid (pNL4-3.Luc.R-S-) containing previously described stabilizing mutations in the capsid protein (PMID: 12547912) [43] and firefly luciferase in the *nef* open reading frame (1.35ug) and pSARS-CoV-2 expressing SARS-CoV-2 S protein (0.6ug) (GenBank; MN908947.3), a gift from Paul Bieniasz [41, 44]. Transfection was performed in 293T/17 cells using genejuice (Novagen, USA) transfection kit according to manufacturer’s protocol. At day 3 or day 4, pseudotyped SARS-CoV-2 virus particles were harvested and filtered over a 0.45 µm nitrocellulose membrane (Sartorius Stedim, Gottingen, Germany). SARS-CoV-2 pseudovirus productions were quantified by RETRO-TEK HIV-1 p24 ELISA according to manufacturer instructions (ZeptoMetrix Corporation).

### SARS-CoV-2 isolate (hCoV-19/Italy-WT)

The wild-type (WT) authentic SARS-CoV-2 virus hCoV-19/WT (D614G variant) was obtained from Dr. Maria R. Capobianchi through BEI Resources, NIAID, NIH: SARS-Related Coronavirus 2, Isolate Italy-INMI1, NR-52284, originally isolated on January 2020 in Rome, Italy. SARS-CoV-2 authentic virus stocks from primary isolates were generated in VeroE6 cells. Cytopathic effect (CPE) formation was monitored and after 48h the virus supernatant was harvested. Viral titers were determined by tissue cultured infectious dose (TCID50) on VeroE6 cells. Briefly, VeroE6 cells were seeded in a 96 well-plate at a cell density of 10000 cells/well. The following day, cells were inoculated with a 5-fold serial dilution of SARS-CoV-2 isolate in quadruplicate. Cell cytotoxicity was measured by using an MTT assay 48h post infection. Loss of MTT staining, analyzed by spectrometer (OD_580nm_) was indicative of SARS-CoV-2 induced CPE. Viral titer was determined as TCID50/mL and calculated based on the method first proposed by Reed & Muench [45]. All experiments with the WT SARS-CoV-2 isolate (hCoV-19/Italy-WT) were performed in a BSL-3 laboratory, following appropriate safety and security protocols approved by the Amsterdam UMC BioSafetyGroep and performed under the environmental license obtained from the municipality of Amsterdam.

### SARS-CoV-2 (hCoV-19/Italy-WT) neutralization assay

Antibody neutralization activity of SARS-CoV-2 infection was tested as previously described [46] and including some modifications. Briefly, VeroE6 cells were seeded at a density of 10,000 cells/well in a 96-well plate one day prior to the start of the neutralization assay. Heat inactivated sera samples were serially diluted in cell culture medium CO_2_ independent medium (Gibco life Technologies, Gaithersburg, Md.) supplemented with 10% fetal calf serum (FCS), L-glutamine and penicillin/streptomycin (10µg/mL), mixed 1:1 ratio with authentic SARS-CoV-2 and incubated for 1h at 37°C. Subsequently, these mixtures were added to the cells in a 1:1 ratio and incubated for 48h at 37°C without CO_2,_ followed by an MTT assay. Loss of MTT staining, analyzed by spectrometer (OD_580nm_) was indicative of SARS-CoV-2 induced CPE. The neutralization titers (IC50) were determined as the serum dilution or antibody concentration at which infectivity was inhibited by 50%, respectively, using a non-linear regression curve fit (GraphPad Prism software version 8.3) and serum dilutions were converted into international units per mL (IU/mL) using the WHO International Standard for anti-SARS-CoV-2 immunoglobulin (NIBSC 20/136).

### Opsonization assay of SARS-CoV-2 pseudovirus and hCoV-19/Italy-WT

Incubation of SARS-CoV-2 with pre-COVID-19 pandemic pooled normal human serum (NHS) mediated covalent deposition of C3 fragments (C3b, iC3b,C3d, C3c) and specific-IgGs on the viral surfaces To mimic the *in vivo* situation [22], where SARS-CoV-2 is opsonized with complement or IgGs, SARS-CoV-2 pseudovirus (191.05 ng/mL of SARS-CoV-2 pseudovirus) and authentic SARS-CoV-2 (hCoV-19/Italy-WT, 1000TCID/mL) were incubated for 1 h at 37°C with pre-pandemic NHS (1:10 ratio), as complement source to obtain complement-opsonized SARS-CoV-2; with specific IgGs or highly IgG content sera, to obtains IgG-opsonized SARS-CoV-2, or a combination of both (C-Ig) in a 1:10 ratio [47]. As negative control, the virus was incubated under same conditions with plain RPMI1640. Serum from at least 10 healthy donors, referred to as NHS, were pooled and stored at -80°C. The presence of C3 fragments and IgGs on the viral surface was detected by opsonization ELISA assay, also called Viral Capture Assay (VCA) as previously described [48–50]. Briefly, 96-well plates were coated with rabbit anti-mouse IgG (DAKO) at 4°C overnight. ELISA plates were coated anti-human C3c and C3d as well as human IgG and incubated overnight with differentially opsonized virus preparations (1ng/p24 per well) at 4°C and extensively washed with RPMI1640 medium to remove unbound virus. Mouse IgG antibodies was used as control for background binding. For the SARS-CoV-2 pseudovirus, viral samples were lysed (1% Triton) and binding was quantified by p24 ELISA to confirm the opsonization pattern [51]. The opsonization pattern of SARS-CoV-2 isolate (hCoV-19/Italy-WT) was determined by qPCR. The viral samples were lysed, and SARS-CoV-2 RNA was extracted using FavorPrep Viral RNA Minikit (FAVORGEN, Ping-Tung, Taiwan), according to the manufacturer’s instructions. Sequences specific to region N1 of the Nucleocapsid gene published on the CDC website (https://www.cdc.gov/coronavirus/2019-ncov/lab/rt-pcr-panel-primer-probes.html) were used. Luna Universal Probe One-Step RT-PCR kit (New England BioLabs, Ipswich, Mass) was used for target amplification, and runs were performed on the CFX96 real-time detection system (Bio-Rad). For absolute quantification using the standard curve method, SARS-CoV-2 RNA was obtained as a PCR standard control from the National Institute for Biological Standards and Control UK (Ridge, UK).

### COVID-19 patient serum

Pooled serum from at least 20 random individuals (mild/moderate disease) all in 2020 (Wuhan variant, no vaccination), collected 19 days post-symptom onset with high neutralizing antibody content, was heat inactivated (1 h at 56°C) to destroy complement activity. In brief, SARS-CoV-2 isolate (hCoV-19/Italy-WT, 1000TCID/mL) was incubated for 1 h at 37°C with high neutralizing antibody pooled serum in a 1:10 ratio, to generate antibody-opsonized SARS-CoV-2.

Serum from 10 COVID-19 patients after natural infection with either mild (5x) or severe (5x) disease outcome, collected approximately 3 months post-infection, were used in a 1:10 ratio as antibody mediated-complement opsonization source for opsonization and generate complement- and antibody-opsonized SARS-CoV-2. The ability to opsonize the virions was assessed by ELISA, as previously described [48, 49].

### Virus binding and internalization

In order to determine SARS-CoV-2 binding and internalization, target cells were seeded in a 96-well plate at a density of 100,000 cells in 100µl. Cells were exposed to SARS-CoV-2 isolate (hCoV-19/Italy-WT, 1000TCID/mL) for 4 h at 4°C for binding and as well 4 h at 37°C for internalization. After 4 h incubation, cells were washed extensively to remove the unbound virus. Cells were lysed with AVL buffer and RNA was isolated with the QIAmp Viral RNA Mini Kit (Qiagen) according to the manufacturer’s protocol.

### DC Stimulation and infection

DCs were left unstimulated or stimulated with 10ng/mL LPS *Salmonella typhosa* (Sigma) and SARS-CoV-2 isolate (hCoV-19/Italy-WT) with different opsonization patterns SARS-CoV-2, SARS-CoV-2-C, SARS-CoV-2-Ab and SARS-CoV-2-Ab/C at 1000TCID/mL.

Blocking of CR3/CD11b (LEAF-purified CR3/CD11b) and CR4/CD11c (LEAF-purified CR4/CD11c) was performed with 10µg/mL for 30 min at 37°C before adding the virus preparations. Similarly, for blocking of FcγRII, DCs were treated with anti-CD32 antibody 1µg/mL for 1 h at 37°C. DCs do not express ACE2 and are therefore not infected by SARS-CoV-2 [33]. Therefore, viral stimulation SARS-CoV-2 isolate (hCoV-19/Italy-WT) at 1000TCID/mL (MOI 0.028) was performed for 2 h, 6 h, 8 h (not shown) after which the cells were lysed for RNA isolation and cytokine production analysis. In addition, cells were stimulated for 24 h and fixed for 30 min with 4% paraformaldehyde to assess the maturation phenotype with flow cytometry.

### Caspase-1 activity

Active caspase-1 was detected using the FAM-FLICA Caspase-1 Assay kit (Immunochemistry Technologies) according to manufacturer’s instructions. In brief, DCs were washed in IMDM medium lacking phenol red (Gibco), supplemented with 10% FCS, penicillin and streptomycin (100 U/mL and 100 μg/mL, respectively, Thermo Fisher) prior to stimulations. After 14 h, DCs were treated with FAM FLICA reagent and incubated at 37°C, 5% CO2 for 1 h. Cells were washed three times in apoptotic wash buffer (Immunochemistry Technologies), immediately followed by flow cytometry analysis using the FACSCanto II (BD Bioscience) and FlowJo software version 10.7 and guidelines for the use of flow cytometry and cell sorting in immunological studies were followed. Live cells were gated based on FSC/SSC and the caspase-1^+^ (FAM-FLICA) population within this live cell population was assessed.

### RNA isolation and quantitative real-time PCR

Cells incubated with SARS-CoV-2 isolate (hCoV-19/Italy-WT) were lysed with AVL buffer and RNA was isolated with QIAmp Viral RNA Mini Kit (Quiagen) according to the manufactureŕs protocol. cDNA was synthesized with M-MLV reverse transcriptase Kit (Promega) and diluted 1/5 before further application. PCR amplification was performed by using RT-PCR in the presence of SYBR green in a 7500 Fast Realtime PCR System (ABI). Specific primers were designed with Primer Express 2.0 (Applied Biosystems). The following primer sequences were used:

GAPDH: F-primer 5’-CCATGTTCGTCATGGGTGTG-3’, R-primer 5’-GGTGCTAA GCAGTTGGTGGTG-3’; SARS-CoV-2 ORF1b: F-primer 5’- TGGGGTTTTACAGGTAACCT-3’, R-primer 5’-AACACGCTTAACAAAGCACTC-3’; IFNB: F-primer 5’-ACAGACTTACAGGTTACCTCCGAAAC-3’, R-primer 5’- CATCTGCTGGTTGAAGAATGCTT-3’; CXCL10: F-primer 5’- CGCTGTACCTGCATCAGCAT-3’; R-primer 5’-CATCTCTTCTCACCCTTCTTTTTCA-3’; IL-6: F-primer 5’-TGCAATAACCACCCCTGACC-3’, R-primer 5’- TGCGCAGAATGAGATGAGTTG-3’; IL-10: F-primer 5’-GAGGCTACGGCGCTGTCAT- 3’, R-primer 5’-CCACGGCCTTGCTCTTGTT-3’; IL-12p35: F-primer TGGACCACCTCAGTTTGGC; R-primer TTCCTGGGTCTGGAGTGGC;

The normalized amount of target mRNA was calculated from the Ct values obtained for both target and household mRNA with the equation Nt = ^2Ct^ ^(GAPDH)−Ct(target)^.

### ELISA

Cell supernatants were harvested after 24 h of stimulation and secretion of IL-1β was measured by ELISA (eBiosciences) according to the manufacturer’s instructions. Supernatant containing SARS-CoV-2 was inactivated with 1% triton before performing the ELISA. OD_450nm_ values were measured using BioTek Synergy HT.

### Statistics

All results are presented as mean ± SEM and were analyzed by GraphPad Prism 9 software (GraphPad Software Inc.). A two-tailed, parametric Student’s *t*-test for paired observations (differences within the same donor) or unpaired observation, Mann-Whitney tests (differences between different donors, that were not normally distributed) was performed. For unpaired, non-parametric observations a one-way ANOVA or two-way ANOVA test with post hoc analysis (Tukey’s or Dunnett’s) were performed. Statistical significance was set at *P< 0.05, **P<0.01***P<0.001****P<0.0001. The allele and genotype frequencies were determined by direct counting. The significant differences of allele genotype frequencies were calculates using Pearson χ2 test.

## Results

### Complement-opsonized SARS-CoV-2 (hCoV-19/Italy-WT) activates dendritic cells via CR3/CD11b and CR4/CD11c

DCs neither become infected nor activated by SARS-CoV-2 (hCoV-19/Italy-WT) [33, 52]. Here we investigated whether complement-opsonized SARS-CoV-2 interacts with DCs. Incubation of pseudotyped SARS-CoV-2 and SARS-CoV-2 isolate (hCoV-19/Italy-WT) with pre-COVID-19 pandemic normal human serum (NHS), containing no virus-specific antibodies, led to efficient opsonization of SARS-CoV-2 as observed by detection of C3c and C3d, but as expected not IgGs were found on the virus surface. This was determined by ELISA for complement proteins and antibodies [49, 51] and qPCR (Fig. 1A, Supp 1A). Notably, complement-opsonized SARS-CoV-2 isolate (hCoV-19/Italy-WT) bound more strongly to DCs than non-opsonized SARS-CoV-2 (Fig. 1B). Blocking antibodies against the α-chain of complement receptors CR3/CD11b and CR4/CD11c abrogated complement-opsonized SARS-CoV-2 binding to DCs.

**Figure 1.**
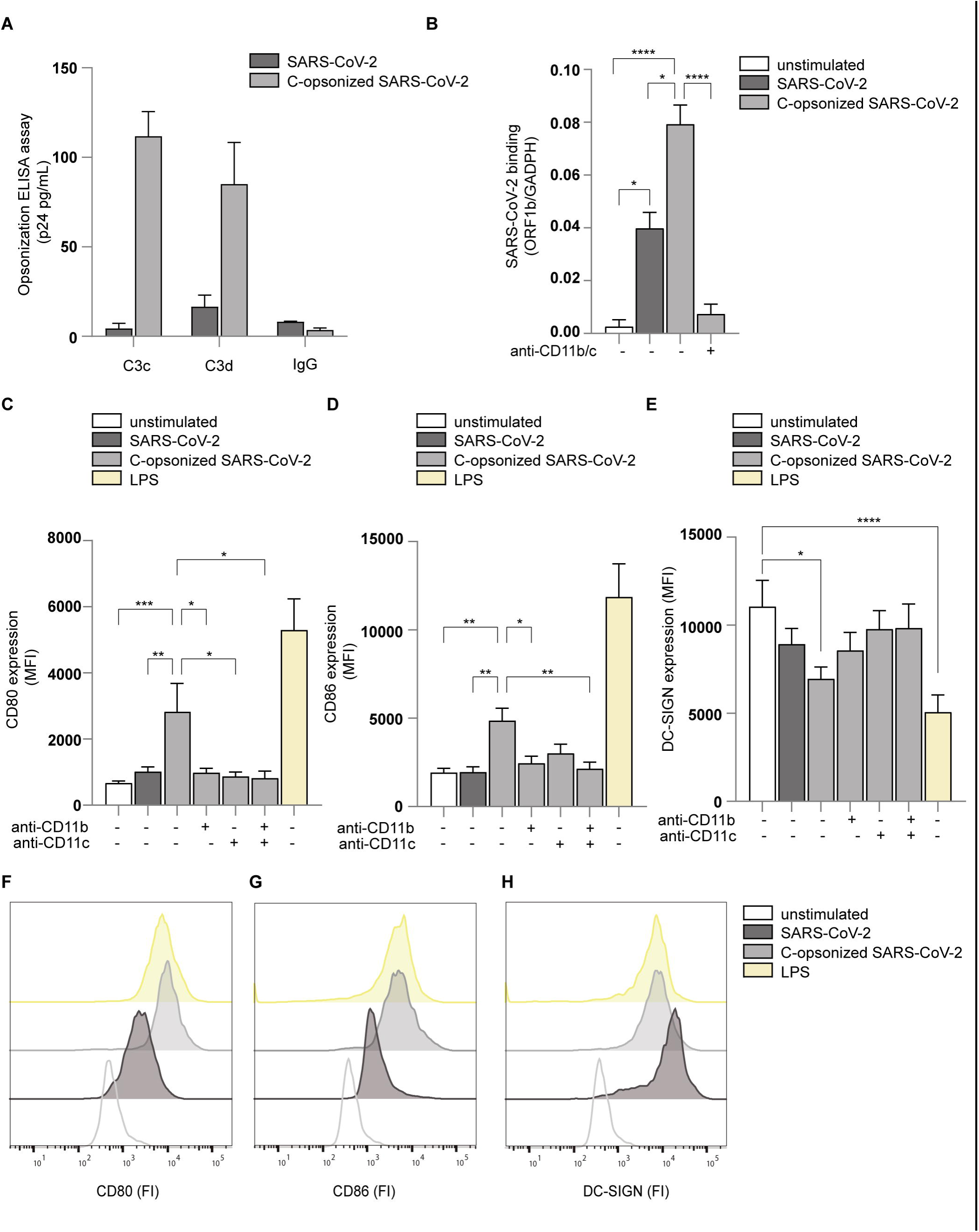
Complement-opsonized SARS-CoV-2 activates DCs via CD11b and CD11c. (A) SARS-CoV-2 pseudovirus opsonization by C3c and C3d was determined by ELISA (p24 pg/mL) (n=3). (B) Human monocyte-derived DCs were exposed to SARS-CoV-2 isolate (hCoV-19/Italy-WT, 1000TCID/mL) and complement-opsonized SARS-CoV-2 (hCoV-19/Italy-WT, 1000TCID/mL) for 4 h in presence or absence of antibodies against CD11b and CD11c. Virus binding was determined by quantitative real-time PCR (n=6 donors). (C-E) DCs were exposed to SARS-CoV-2 or complement opsonized SARS-CoV-2 for 24 h and expression of CD80, CD86 and DC-SIGN was determined by flow cytometry (n=12 donors). LPS stimulation was used as positive control. (F-H) Representative histograms of CD80 (F), CD86 (G) and DC-SIGN (H) expression. Data show the mean values and error bars are the SEM. Statistical analysis was performed using (B) ordinary one-way ANOVA with Tukey multiple-comparison test. *p ≤ 0.05, ****p ≤ 0.0001(n=6 donors). (C-E) 2-way ANOVA with Tukey multiple-comparison test. *p ≤ 0.05, **p ≤ 0.01, ***p ≤ 0.001, ****p ≤ 0.0001 (n=12 donors).

We next investigated induction of DC maturation by complement-opsonized SARS-CoV-2 isolate (hCoV-19/Italy-WT) by analysing expression of co-stimulatory molecules CD80 and CD86, DC-SIGN, and complement receptors CR3/CD11b and CR4/CD11c. In contrast to SARS-CoV-2, complement-opsonized SARS-CoV-2 induced significant expression of CD80 (Fig. 1C, F) and CD86 (Fig. 1D, G) to similar levels as observed for LPS. Upregulation of CD80 and CD86 was abrogated by blocking antibodies against CR3/CD11b and CR4/CD11c alone or in combination (Fig. 1C, D). DC-SIGN expression was reduced by complement-opsonized SARS-CoV-2 (Fig. 1E, H), similar as observed for LPS, and expression was restored in presence of blocking antibodies against CR3/CD11b and/or CR4/CD11c (Fig. 1E). These results strongly suggest that complement-opsonization enhances SARS-CoV-2 capture by DCs and induces DC maturation via CR3 and CR4 in contrast to non-opsonized SARS-CoV-2.

### Complement-opsonized SARS-CoV-2 (hCoV-19/Italy-WT) induces type I IFN and cytokine responses

Next, we investigated whether complement-opsonized SARS-CoV-2 isolate (hCoV-19/Italy-WT) induces antiviral type I interferon (IFN) as well as cytokine responses. Notably, in contrast to non-opsonized SARS-CoV-2, complement-opsonized SARS-CoV-2 isolate (hCoV-19/Italy-WT) induced significantly higher mRNA levels of IFNβ as well as IFN-stimulated genes (ISGs) APOBEC3G, IRF7 and CXCL-10 (Fig. 2A-D). Blocking antibodies against CR3/CD11b and CR4/CD11c abrogated the induction of IFNβ and ISGs to similar levels as observed with SARS-CoV-2 alone (Fig.2A-D). These data strongly suggest that, in contrast to its non-opsonized counterpart, complement-opsonized SARS-CoV-2 induces antiviral type I IFN responses via CR3 and CR4.

**Figure 2.**
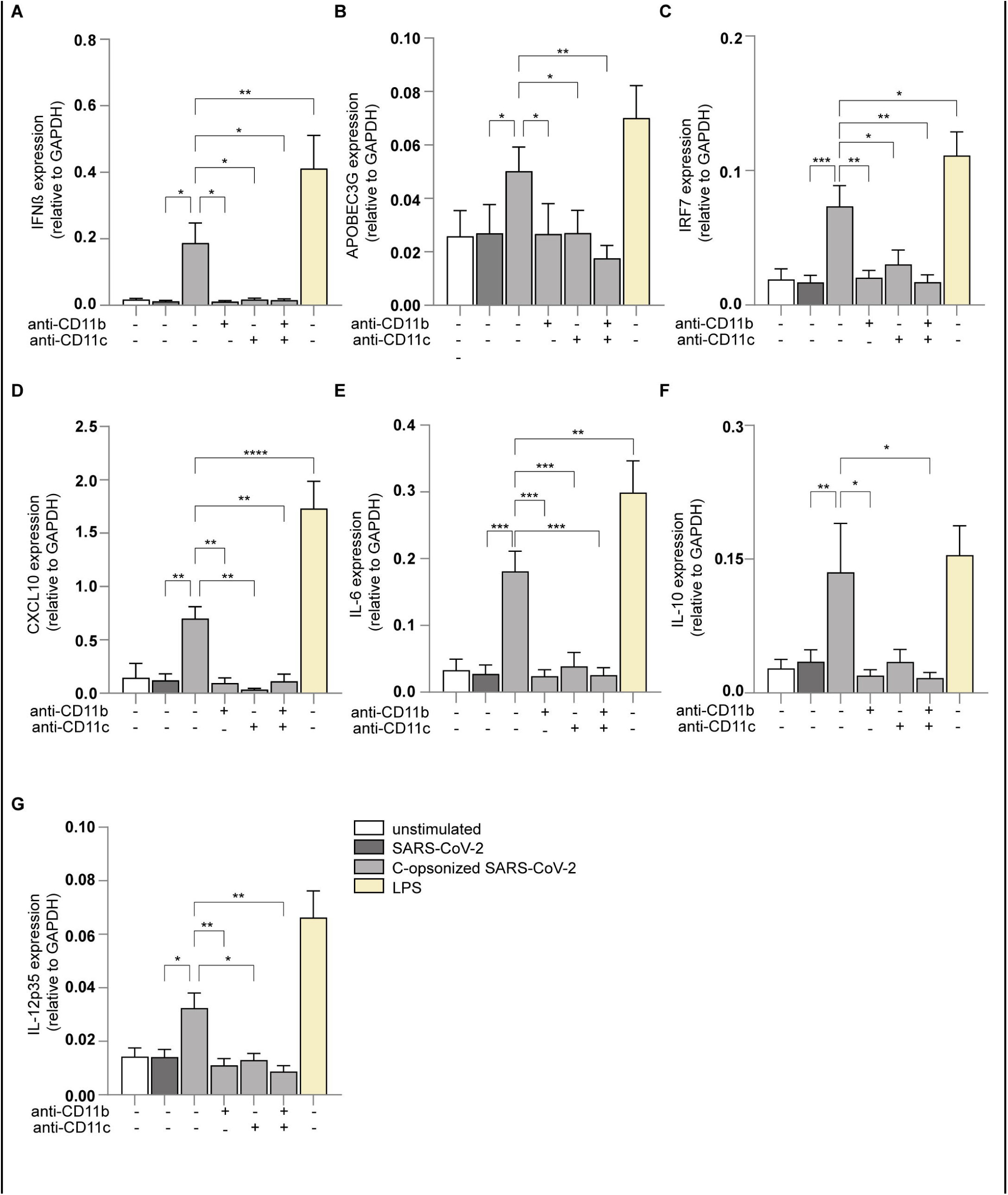
Complement-opsonized SARS-CoV-2 induces type I IFN and cytokine responses. (A-G) Human monocyte-derived DC were exposed to SARS-CoV-2 isolate (hCoV-19/Italy-WT, 1000TCID/mL), complement-opsonized SARS-CoV-2 (hCoV-19/Italy-WT, 1000TCID/mL) and LPS (100 ng/mL) in presence or absence of antibodies against CD11b and CD11c for 2 h and 6 h. mRNA levels of IFN-β (A), APOBEC3G (B), IRF7 (C), CXCL10 (D), IL-6 (E), IL-10 (F) and IL-12p35 (G) were determine with qPCR after 2 h (A) and after 6 h (B-G) (n=14 donors). Data show the mean values and error bars are the SEM. Statistical analysis was performed using (A-F) 2-way ANOVA with Dunnett’s multiple-comparison test. *p ≤ 0.05, **p ≤ 0.01, ***p ≤ 0.001, ****p ≤ 0.0001 (n=14 donors).

Moreover, complement-opsonized SARS-CoV-2 induced transcription of cytokines IL-6, IL-10 and IL-12p35, and expression was abrogated by blocking CR3/CD11b and CR4/CD11c (Fig. 2E-G). Secretion of biologically active IL-1β is tightly regulated and depends on induction of pro-IL1β and caspase-dependent processing into IL-1β, which is subsequently secreted by DCs [53–55]. Notably, complement-opsonized SARS-CoV-2 induced secretion of IL-1β protein in contrast to non-opsonized SARS-CoV-2, and IL-1β production was inhibited by antibodies against CR3/CD11b and CR4/CD11c (Fig. 3A). As caspase-1 is an important caspase involved in pro-IL-1β processing, we investigated whether complement-opsonized SARS-CoV-2 activates caspase-1 using the FAM FLICA assay. Complement-opsonized SARS-CoV-2 significantly increased active caspase-1 in DCs (Fig. 3B and Supp. 1C), similar as observed for LPS- and ATP-stimulated DCs (Fig. 3B and Supp. 1D). Caspase-1 activation by complement-opsonized SARS-CoV-2 was blocked by antibodies against CR3/CD11b and CR4/CD11c. Non-opsonized SARS-CoV-2 did not induce caspase-1 activation.

**Figure 3.**
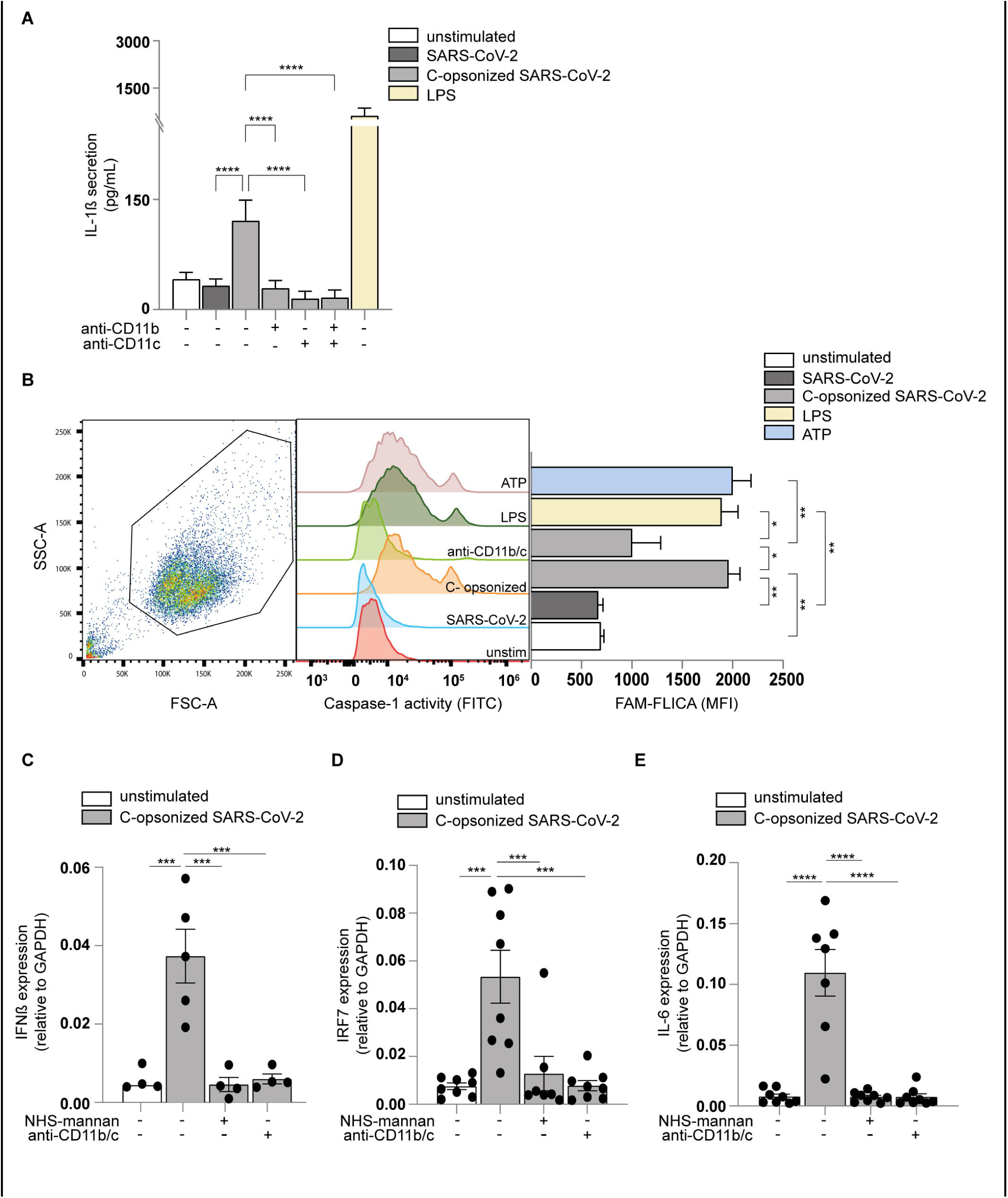
Caspase-1 activity is enhanced by activated DCs by complement-opsonized SARS-CoV-2. (A) Human monocyte-derived DC were exposed to SARS-CoV-2 isolate (hCoV-19/Italy-WT, 1000TCID/mL), complement-opsonized SARS-CoV-2 (hCoV-19/Italy-WT, 1000TCID/mL) and LPS (100 ng/mL) in presence or absence of antibodies against CD11b and CD11c and IL-1β secretion (pg/mL) in the supernatant was measured after 24 h by ELISA (n=8). (B) DCs were left unstimulated or treated with anti-CD11b/c prior exposure to non- and complement-opsonized SARS-CoV-2, LPS and ATP. DCs with active caspase-1 were detected after 14 h by flow cytometry using the FAM-FLICA assay (n=3). (C-E) NHS was incubated with mannan, prior SARS-CoV-2 opsonization. DCs were exposed to non-, complement-opsonized SARS-CoV-2 and NHS-mannan opsonized SARS-CoV-2 in presence or absence of anti-CD11b/c, and mRNA levels of IFNβ (C), IRF7 (D) and IL-6 (E) were determined by qPCR (n=8 donors). Data show the mean values and error bars are the SEM. Statistical analysis was performed using (A) 2-way ANOVA with Tukey multiple-comparison test. ****p ≤ 0.0001 (n=8donors). (B) ordinary one-way with Tukey’s multiple-comparison test. *p ≤ 0.05, **p ≤ 0.01 (n=3 donors). (C-E) 2-way ANOVA with Tukey multiple-comparison test. ***p ≤ 0.001, ****p ≤ 0.0001 (H) (n=8 donors).

Next, we investigated whether the lectin pathway was involved in complement activation by SARS-CoV-2. Pre-treatment of pre-COVID-19 pandemic NHS with mannan, which act as an inhibitor of the lectin pathway via MBL [56], significantly decreased DC-induced type I IFN and IL-6 responses (Fig. 3C-E), indicating that SARS-CoV-2 activates complement by the lectin pathway through the carbohydrate-recognition domain. Together these data strongly suggest that complement-opsonization of SARS-CoV-2 by MBL induces a potent pro-inflammatory as well as an antiviral type I IFN response in DCs via CR3 and CR4.

### Convalescent serum from COVID-19 patients blocks immune responses induced by complement-opsonized SARS-CoV-2 isolate (hCoV-19/Italy-WT) via CD32

Antibodies are important in induction of complement activation and subsequent deposition [57, 58]. Here we investigated whether antibodies against SARS-CoV-2 affect complement-induced immunity by DCs. Serum from 20 (mild/moderate disease) recovered COVID-19 patients neutralized SARS-CoV-2 infection (Supp. 2A), indicating that serum contains neutralizing antibodies against SARS-CoV-2 [41, 46, 59]. Next, we investigated the effect of COVID-19 serum on complement-induced immunity. Complement-opsonized SARS-CoV-2 (hCoV-19/Italy-WT) induced ISGs APOBEC3G, IRF7 and CXCL10 as well as cytokines IL-6, IL-10 and IL-12p35 (Fig. 4A-F). Notably, pre-incubation of complement-opsonized SARS-CoV-2 with COVID-19 convalescent serum abrogated type I IFN as well as cytokine responses. The complement-induced responses were restored by blocking FcγRII (CD32) using an inhibitory CD32 antibody (Fig. 4A-F). These data strongly suggest that antibodies against SARS-CoV-2 suppress immune responses induced by complement-opsonized SARS-CoV-2 via CD32.

**Figure 4.**
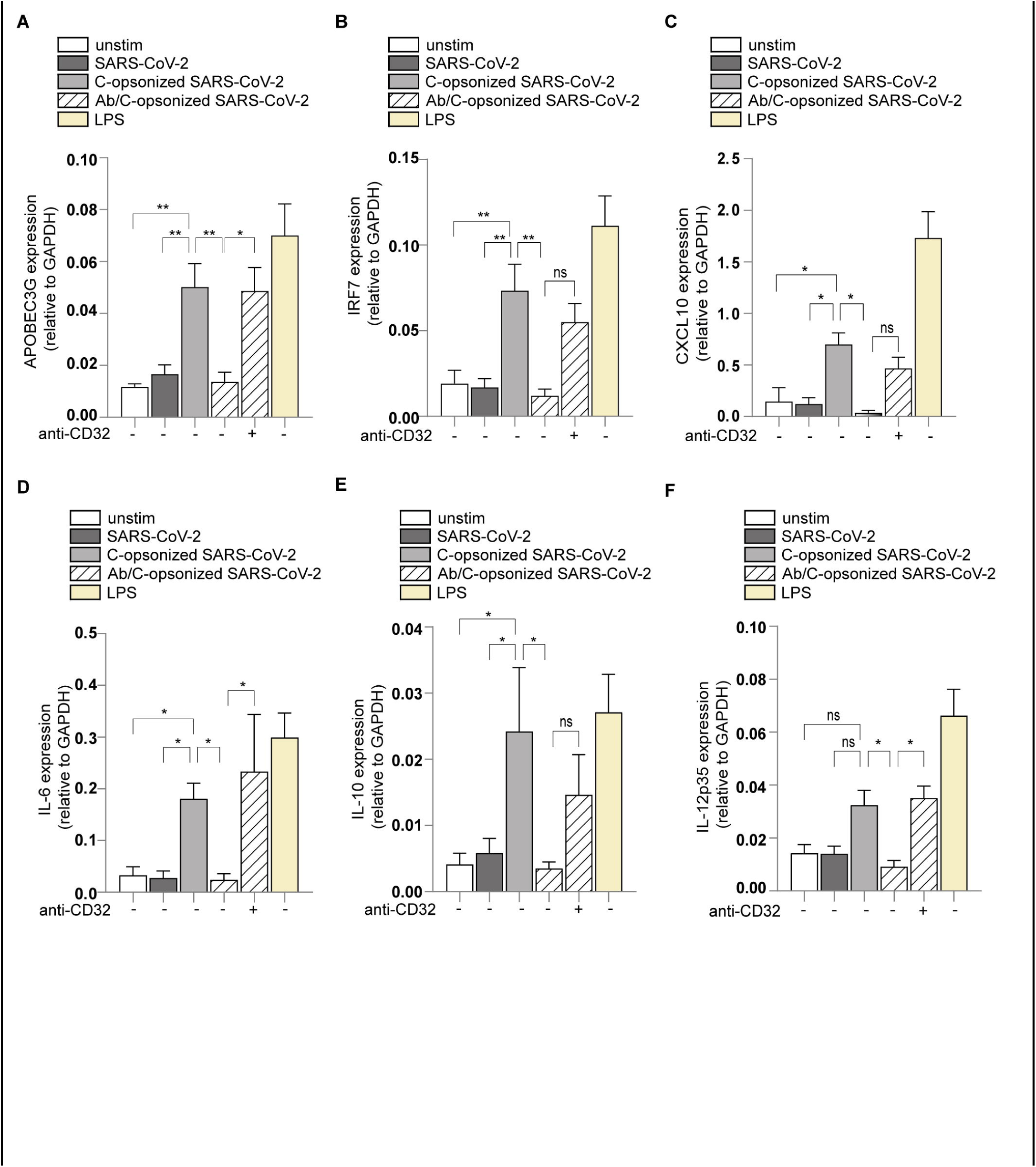
anti-SARS-CoV-2 antibodies present in sera suppress complement activation mediated immune activation via CD32. (A-F) Human monocyte-derived DCs were exposed to SARS-CoV-2 isolate (hCoV-19/Italy-WT, 1000TCID/mL), complement-opsonized SARS-CoV-2 (hCoV-19/Italy-WT, 1000TCID/mL) and LPS (100 ng/mL) in presence or absence of anti-CD32 for 6 h. mRNA levels of APOBEC3G (A), IRF7 (B), CXCL10 (C), IL-6 (D), IL-10 (E) and IL-12p35 (F) were determine with qPCR after 6h (n=14 donors). Data show the mean values and error bars are the SEM. Statistical analysis was performed using (A-F) 2-way ANOVA with Dunnett’s multiple-comparison test. *p ≤ 0.05, **p ≤ 0.01, ***p ≤ 0.001, ****p ≤ 0.0001 (n=14 donors).

### Monoclonal antibodies against SARS-CoV-2 block complement-induced immunity to SARS-CoV-2

We investigated whether monoclonal antibodies against SARS-CoV-2 suppress the inflammation induced by complement-opsonized SARS-CoV-2 and how this is affected by the neutralizing capacity. We compared the effect of non-neutralizing and neutralizing antibodies against SARS-CoV-2 isolated from COVID-19 patients, COVA1-27 and COVA1-18, respectively [41]. Complement-opsonized SARS-CoV-2 induced IFNβ transcription and, notably, pre-incubation of complement-opsonized SARS-CoV-2 with either neutralizing (COVA1-18) or non-neutralizing antibody (COVA1-27) against SARS-CoV-2 abrogated IFNβ transcription (Fig. 5A). The COVA1-27-mediated suppression was abrogated by CD32 inhibition thereby restoring IFNβ transcription to levels observed with complement-opsonized SARS-CoV-2 (Fig. 5A). CD32 inhibition had less effect on COVA1-18-mediated suppression (Fig. 5A). Similarly, both COVA1-18 and 1-27 suppressed induction of ISGs APOBEC3G, IRF7 and CXCL10 as well as cytokines IL-6 and IL-10 (Fig. 5B-F). CD32 inhibition restored induction of CXCL10 and partially for APOBEC3G and IRF7. In contrast to IL-6, IL-10 induction was restored by CD32 inhibition. Moreover, induction of CD86 was also suppressed by COVA1-18 and 1-27, which was restored by CD32 inhibition (Supp. 2B). These data strongly suggest that antibodies against SARS-CoV-2 present in serum inhibit complement-induced immune responses via CD32 and this is independent of neutralization capacity.

**Figure 5.**
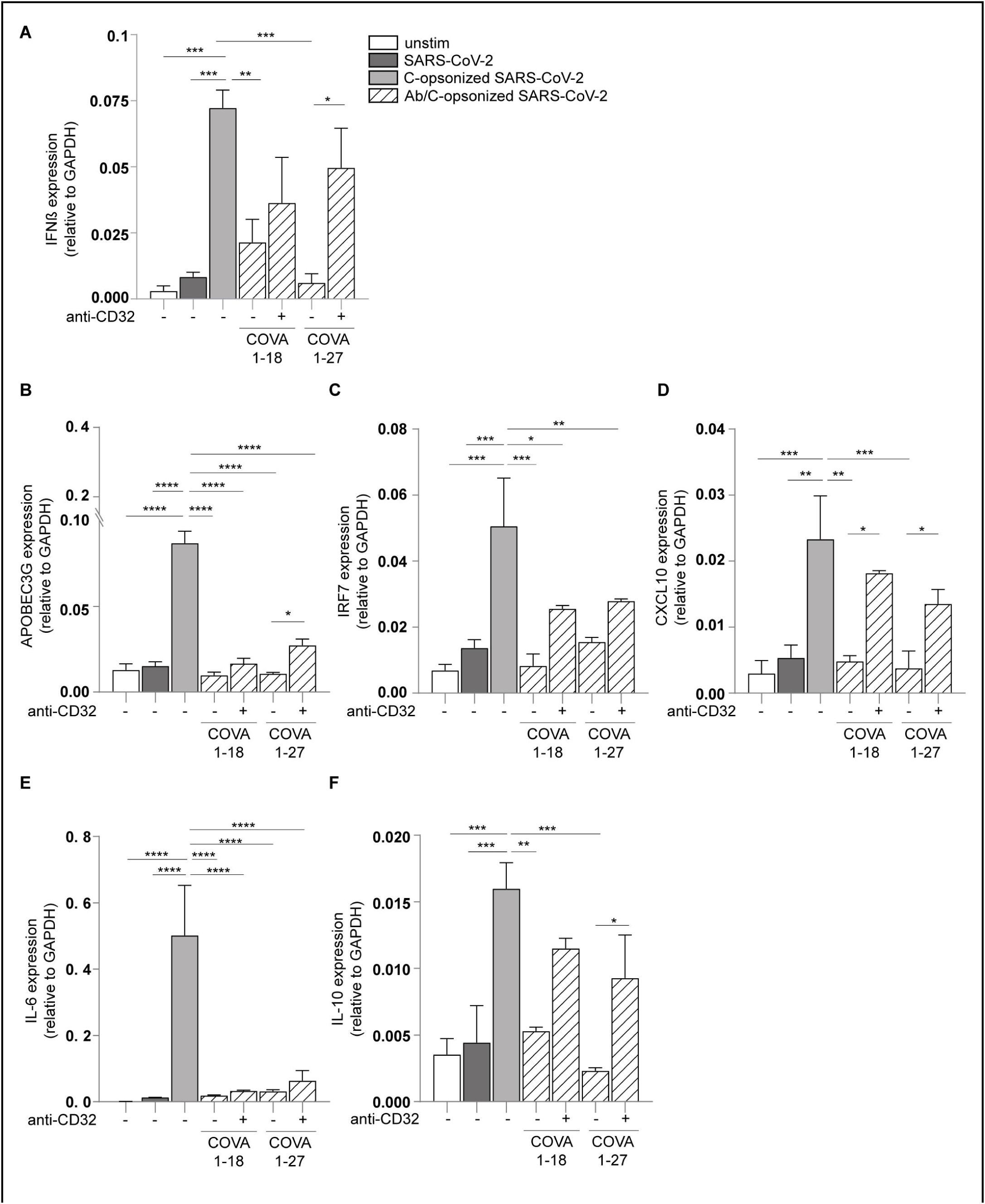
non- and neutralizing anti-SARS-CoV-2 antibodies suppress complement activation mediated immune activation via CD32. (A-F) SARS-CoV-2 was pre-incubated with patient isolated mAb COVA1-18 and COVA1-27 (10µg/mL) for 30 min at 37°C. Human monocyte-derived DCs were exposed to SARS-CoV-2 isolate (hCoV-19/Italy-WT, 1000TCID/mL) alone or with blocks, to complement-opsonized SARS-CoV-2 (hCoV-19/Italy-WT, 1000TCID/mL) and LPS (100 ng/mL) in presence or absence of anti-CD32 for 2 h and 6 h. mRNA levels of IFNβ (A), APOBEC3G (B), IRF7 (C), CXCL10 (D), IL-6 (E) and IL-10 (F) were determined by qPCR (n=6 donors (A) and (n=4 donors) (B-F). Data show the mean values and error bars are the SEM. Statistical analysis was performed using (A-F) 2-way ANOVA with Tukey’s multiple-comparison test. *p ≤ 0.05, **p ≤ 0.01, ***p ≤ 0.001, ****p ≤ 0.0001, (A) (n=6donors) and (B-F) (n=4 donors).

### Serum samples from mild and severe COVID-19 patients block complement-induced immunity to SARS-CoV-2

Circulating immune complexes have been correlated with complement activation in severe/critical COVID-19 patients [59–62]. To analyse the impact of antibody status and complement function in parallel, we screened serum from 5 individuals with mild outcome *versus* 5 severe COVID-19. Serum from both groups were incubated for 1 h at 37°C with SARS-CoV-2 isolate (hCoV-19/Italy-WT) and the presence of C3c/C3d and IgGs was determined by ELISA (Fig. 6A). Opsonization of SARS-CoV-2 with serum from either mild or severe COVID-19 patients led to the deposition of C3c and C3d fragments as well as IgG on SARS-CoV-2. Serum from mild COVID-19 patients caused inferior opsonization by C3c/d compared to IgG, whereas serum from severe COVID-19 patients induced more C3c/d opsonisation compared to IgG. These results suggest that in severe COVID-19 patients, complement is fully activated. DC from healthy donors were stimulated with opsonized-SARS-CoV-2 with serum from either mild or severe COVID-19 patients and induction of IFNβ, ISGs, such as IRF7, and IL-6 was measured. (Fig. 6B-D). Serum from both COVID-19 mild and severe patients suppressed IFNβ, IRF7 and IL-6 induction, which was restored by CD32 inhibition (Fig. 6B-D). Interestingly, CD32 inhibition enhanced both type I IFN and IL-6 responses compared to complement induction alone, which might be due to higher complement concentrations in serum from severe COVID-19 patients as suggested by higher opsonization of SARS-CoV-2 (Fig. 6A). These data suggest that induction of antibodies during COVID-19 disease are important to resolve inflammation.

**Figure 6.**
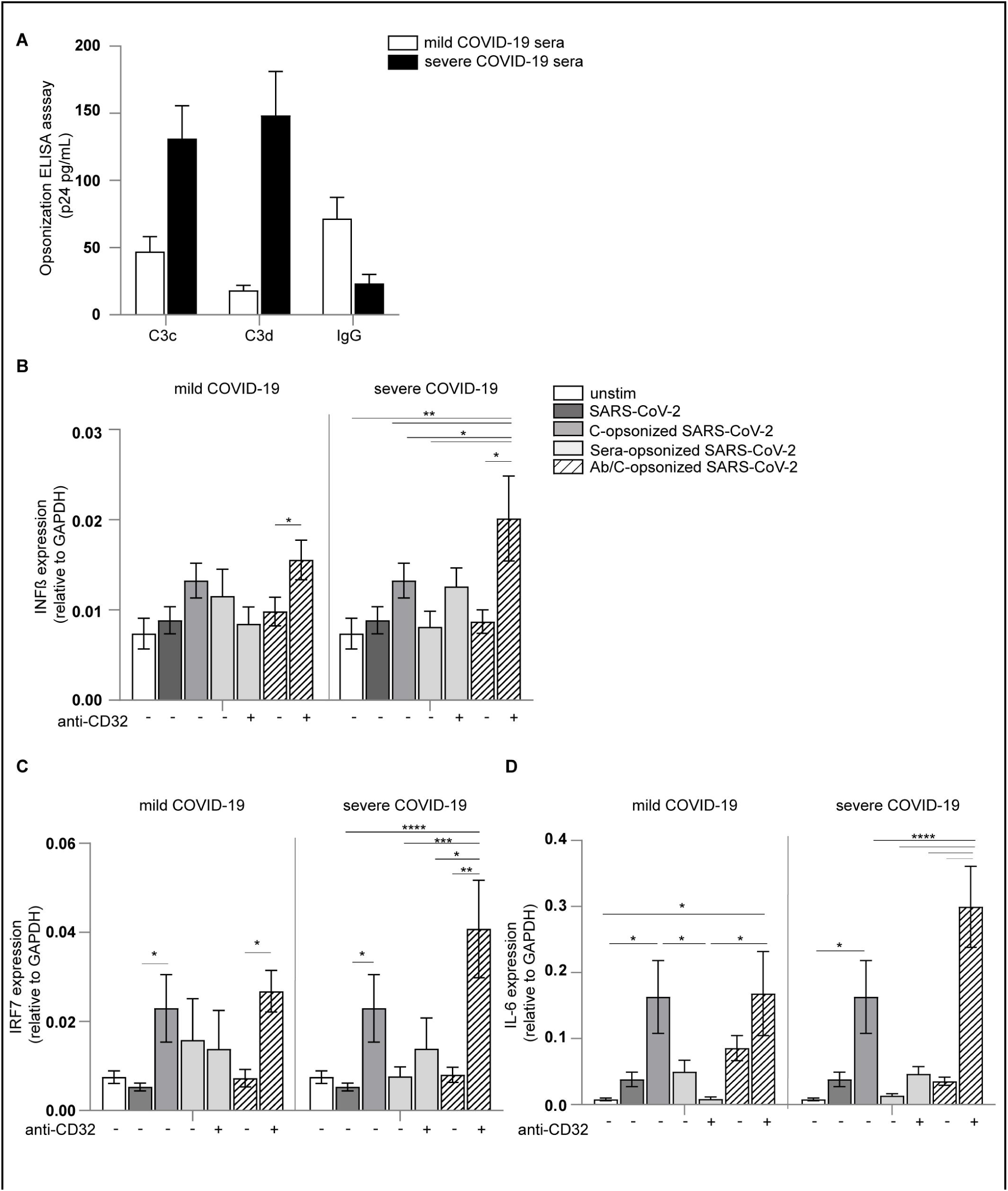
Disease severity dictates SARS-CoV-2 complement activation and antibody response. (A) SARS-CoV-2 pseudovirus opsonization patterns with mild and severe COVID-19 patient sera was determined by ELISA (p24 pg/mL) using anti-human C3c and C3d, for iC3b recognition, and anti-human IgG, for immunoglobulins detection. (B) Human monocyte-derived DCs were exposed to SARS-CoV-2 isolate (hCoV-19/Italy-WT, 1000TCID/mL), to complement-opsonized SARS-CoV-2 (hCoV-19/Italy-WT, 1000TCID/mL), COVID-19 patient serum (mild or severe) and antibody/complement-opsonized SARS-CoV-2 (hCoV-19/Italy-WT, 1000TCID/mL) in presence or absence of anti-CD32 for 2 h and 6 h. mRNA levels for IFNβ (B) were determined after 2 h and mRNA levels of IRF7 (C) and IL-6) (D) after 6 h by qPCR (n=6 donors) (B) and (n=8 donors) (C-D). Data show the mean values and error bars are the SEM. Statistical analysis was performed using (B-D) ordinary one-way ANOVA with Tukey’s multiple-comparison test. *p ≤ 0.05, **p ≤ 0.01 (B) (n=6 donors) and (C-D) (n=8 donors).

## Discussion

Complement is crucial for the induction of inflammatory responses to pathogens leading to an effective adaptive immune response. A hallmark of severe COVID-19 disease is excessive inflammation associated with enhanced morbidity and mortality. Accumulating evidence suggests that overactivation of the complement system contributes to pathophysiology of severe COVID-19 disease. Previously, we have shown that authentic SARS-CoV-2 isolate (hCoV-19/Italy-WT) (1000TCID/mL or MOI 0.028) do not activate DCs, which suggests immune escape. Here we show that SARS-CoV-2 isolate viruses are well opsonized by complement C3b/c fragments and complement-opsonized SARS-CoV-2 efficiently induced DC activation, type I IFN and cytokine responses via complement receptors CR3/CD11b and CR4/CD11c. Notably, we identified antibody responses against SARS-CoV-2 as a negative feedback mechanism in limiting complement-induced inflammation via CD32 signalling, as previously illustrated for HIV-1 (Posch et al, 2012). Our data therefore suggest that complement is crucial in the induction of antiviral innate and adaptive immune responses to SARS-CoV-2 and subsequently elicited antibodies against SARS-CoV-2 downregulate complement-induced immunity thereby preventing aberrant inflammation. This study highlights an important role for antibodies against SARS-CoV-2 to control immune homeostasis and suggest that dysregulation in this control might underlie aberrant inflammatory responses observed in severe COVID-19.

SARS-CoV-2 itself can activate the complement system directly through the lectin pathway [17, 19, 63–65] and the alternative pathway [23]. Specifically, SARS-CoV-2 Spike and nucleocapsid proteins are directly recognized by the lectin pathway components, leading to complement activation and the subsequent complement deposition (C3b) on virions [17]. We observed that inoculation of SARS-CoV-2 isolate (hCoV-19/Italy-WT) with human pre-COVID-19 pandemic serum led to efficient opsonisation of SARS-CoV-2 by C3c and C3d fragments. Opsonization was inhibited by carbohydrate mannan, strongly supporting a role for MBL and the lectin pathway in activating complement. The use of pre-COVID-19 pandemic serum excluded a potential role for antibodies against SARS-CoV-2 in complement activation as has been shown others [22, 28, 66, 67].

SARS-CoV-2 (hCoV-19/Italy-WT) binds to human DCs via heparan sulfate proteoglycans and C-type lectin receptor DC-SIGN [52, 68–70]. SARS-CoV-2 binding to DCs is important for viral transmission to epithelial cells but does not cause immune activation [30, 52, 71]. The lack of immune activation by SARS-CoV-2 is at least partially due to finding that human DCs do not become infected by SARS-CoV-2 as these immune cells do not express ACE2 [30, 72]. Complement-opsonized SARS-CoV-2 was more efficiently bound by DCs than non-opsonized SARS-CoV-2, and binding was inhibited by antibodies against complement receptors CR3/CD11b and CR4/CD11c. In contrast to SARS-CoV-2 alone, complement-opsonized SARS-CoV-2 strongly induced DC maturation as determined by upregulation of co-stimulatory molecules CD80 and CD86. Moreover, complement-opsonized SARS-CoV-2 induced expression of IFNβ and ISGs IRF7, APOBEC3G and CXCL10 as well as cytokines IL-6, IL-10 and IL-1β. Type I IFN responses are pivotal to antiviral immunity by induction of innate resistance to virus replication but also activating cytotoxic T cell and T helper cell responses to viruses [73–75]. In particular, IL-1β is a very potent pro-inflammatory cytokine activating both innate and adaptive immune responses [76, 77]. Our data suggest that complement-opsonized SARS-CoV-2 binding to DCs via CR3 and CR4 leads to pro-IL-1β expression and subsequent activation of Caspase-1 inflammasome and processing of pro-IL-1β into bioactive IL-1β. Although IL-1β induction is important to induce innate and adaptive immunity, unrestrained expression of IL-1β leads to severe inflammation in different diseases [78–81]. Several studies suggest that IL-1β production is an important factor in inflammatory responses during COVID-19 [80, 82–84] but mechanisms that control IL-1β production or type I IFN responses upon SARS-CoV-2 infection remain unidentified.

Antibodies against SARS-CoV-2 have been suggested to activate complement during infection [22, 85–87]. We neither observed induction of complement opsonization of SARS-CoV-2 isolate (hCoV-19/Italy-WT) by serum from COVID-19 patients nor monoclonal human non- or neutralizing antibodies against SARS-CoV-2. In contrast, we observed that serum from COVID-19 patients as well as monoclonal human antibodies against SARS-CoV-2 attenuated complement-opsonized SARS-CoV-2-induced immune inflammatory response. The observed DC maturation as well as type I IFN and cytokine responses induced by complement-opsonized SARS-CoV-2 was inhibited to levels observed for SARS-CoV-2 by serum from mild and severe COVID-19 patients. Similarly, both neutralizing and non-neutralizing antibodies against SARS-CoV-2 blocked these immune responses induced by complement-opsonized SARS-CoV-2.

Human monocyte-derived DCs express the low affinity immunoglobulin FcyRIIa CD32 which is a receptor for IgG and is involved in DC activation [88, 89]. Notably, our data strongly suggest that antibodies against SARS-CoV-2 suppress complement induced immunity by CD32 as blocking antibodies against CD32 restored immune activation induced by complement-opsonized SARS-CoV-2. We observed that non-heat inactivated serum from mild and severe COVID-19 patients similarly suppressed IFNβ, ISG IRF7 and cytokine IL-6 induced by complement-opsonized SARS-CoV-2. We observed that serum from severe COVID-19 patients had enhanced C3c and C3d deposition on SARS-CoV-2 than virus opsonized by serum from mild COVID-19 patients, whereas antibody deposition was increased with serum from mild COVID-19 patients. Although serum from mild and severe patients similarly suppressed immune responses, we observed that blocking CD32 led to an enhanced inflammatory response in presence of serum from severe patients. These data suggest that increased complement-opsonization of SARS-CoV-2 by serum from severe COVID-19 patients will lead more severe inflammation in the absence of or limited antibody responses.

Our data suggest that complement activation by the MBL pathway is important for the induction of innate and adaptive antiviral immunity to SARS-CoV-2 via CR3 and/or CR4 on human DCs. Complement is present in mucosal tissues and this will lead to rapid activation of immunity upon SARS-CoV-2 infection. Our data have uncovered a striking role for antibodies against SARS-CoV-2 in attenuating the complement-induced inflammatory responses and thereby might be required in resolving inflammation. Our findings support a role for antibodies against SARS-CoV-2 induced by vaccinations and after natural infection, not only in limiting infection but importantly in attenuating inflammation upon SARS-CoV-2 infection. Genetic polymorphisms in CD32 signalling pathways involved in attenuating complement-induced immunity might be responsible for unresolved inflammatory responses observed in severe COVID-19. Polymorphisms in MBL and FcyRII have been associated with susceptibility to or severity of some infectious diseases, such SARS-CoV or influenza [90–92] as well as COVID-19 [18, 63, 93, 94], but whether these affect the complement activation and negative feedback mechanism remains to be investigated. We here provide novel immunologic and mechanistic insights into SARS-CoV-2 infection, where the host can cope with the virus due to efficient cellular and humoral immune response. These findings might be exploited for future therapeutic options to improve antiviral immune responses via triggering not yet considered host mechanisms, i.e complement receptors expressed on immune cells.

## Acknowledgements

We thank Jonne Snitselaar, Yoann Aldon and Judith burger for help with production of antibodies and pseudovirus reagents. In addition, we thank Elke Wynberg and Hugo D. G. van Willigen for their contribution to the The *RECoVERED* Study This research was funded by the Netherlands Organisation for Health Research and Development (ZonMw) together with the Stichting Proefdiervrij (ZonMw MKMD COVID-19 grant nr.114025008to TBHG), and European Research Council (Advanced grant 670424t o TBHG), Amsterdam UMC PhD grant and two COVID-19 grants from the Amsterdam institute for Infection & Immunity (to TBHG, RWS, and MJG). This research was supported by a Work Visit Grant of the Amsterdam institute for Infection and Immunity and by an APART-MINT Fellowship of the Austrian Academy of Sciences at the Institute of Hygiene and medical Microbiology of the University of Innsbruck.

## Author contribution

MB-J conceived and designed experiments. MB-J and LD performed the experiments, MB-J, LD and JvH acquired data and analyzed data. MB-J, LD, JvH and TBHG interpreted data and contributed to scientific discussion. DW, G.J.d.B. MJG, NAK and RWS contributed essential research materials and scientific input. MB-J and TBHG wrote the manuscript with input from all listed authors. TBHG perceived of the original study idea and was involved in all aspects of the study. All authors had access to all the data in this study and approved the final version of the manuscript. The corresponding author vouches for the completeness and accuracy of the data.

## Conflict of interest statement

The authors have declared that no conflict of interest exists.

## Ethics approval statement

This study was performed in accordance with the ethical principles set out in the Declaration of Helsinki and was approved by the institutional review board of the Amsterdam University Medical Centers, location AMC Medical Ethics Committee and Ethics Advisory Body of Sanquin Blood Supply Foundation (Amsterdam, The Netherlands). The RECoVERED study was approved by the medical ethical review board of the Amsterdam University Medical Centers (NL73759.018.20). All participants provided written informed consent.

## Funding

This research was further funded through a ZonMW-NWO grant (Dutch Research Council/ Nederlandse organisatie voor Wetenschappelijk Onderzoek) together with the Stichting Proefdiervrij (ZonMW MKMD COVID-19 grant with project number 114025008) as well as the European Research Council (Advanced grant 670424). LEHvdD was supported by the Netherlands Organization for Scientific Research (NWO) (Grant No. 91717305) and MB-J by the ÖAW-APART MINT (Grant No. 11978).

## Figure legends

**Supplementary Figure 1.:**

(A) SARS-CoV-2 RNA copy numbers/mL in pre-coated C3c, C3d and human IgG wells, were detected through qPCR (n=10 donors). (B-C) Human monocyte-derived DCs were exposed to SARS-CoV-2 isolate (hCoV-19/Italy-WT, 1000TCID/mL and complement-opsonized SARS-CoV-2 (hCoV-19/Italy-WT, 1000TCID/mL) in presence or absence of anti-CD11b and anti-CD11c. LPS stimulation was used as positive control for DC maturation, which was measured after 24 h by flow cytometry. Cumulative flow cytometry data of CD11b and CD11c (n=12 donors). (D) Percentages of FLICA^+^ from different stimulated DC (n=3 donors). Data show the mean values and error bars are the SEM. Statistical analysis was performed using (B-C) 2-way ANOVA with Tukey multiple-comparison test. *p ≤ 0.05, **p ≤ 0.01, ***p ≤ 0.001 (n=12 donors). (D) ordinary one-way with Tukey’s multiple-comparison test (n=3 donors).

**Supplementary Figure 2.:**

(A) Neutralization assay using serum collected from 20 confirmed SARS-CoV-2-infected patients at 19 days post-symptom onset (“p.s.o.”). The IC50 for this serum was 58,16 for the Spike-protein. (B) Human monocyte-derived DCs were exposed to SARS-CoV-2 isolate (hCoV-19/Italy-WT, 1000TCID/mL), to complement-opsonized SARS-CoV-2 (hCoV-19/Italy-WT, 1000TCID/mL), to antibody opsonized SARS-CoV-2 (hCoV-19/Italy-WT, 1000TCID/mL) and to antibody/complement-opsonized SARS-CoV-2 (hCoV-19/Italy-WT, 1000TCID/mL) in presence or absence of anti-CD32 for 24 h. LPS stimulation was used as positive control for DC maturation, which was measured after 24 h by flow cytometry. Cumulative flow cytometry data of CD86 (n=12 donors). (C) DCs were stained with antibodies against the surface markers CD16, CD32 and CD64 and analyzed by flow cytometry. Representative histograms for an experiment repeated more than three times with similar results (n=3 donors). Data show the mean values and error bars are the SEM. Statistical analysis was performed using (B) ordinary one-way ANOVA with Tukey multiple-comparison test. *p ≤ 0.05 (n=6 donors).

## Notes

### Competing Interest Statement

The authors have declared no competing interest.

