## Supplementary figures and images for "Antibodies against SARS-CoV-2 control complement-induced inflammatory responses to SARS-CoV-2"

### Suppl. Fig 1

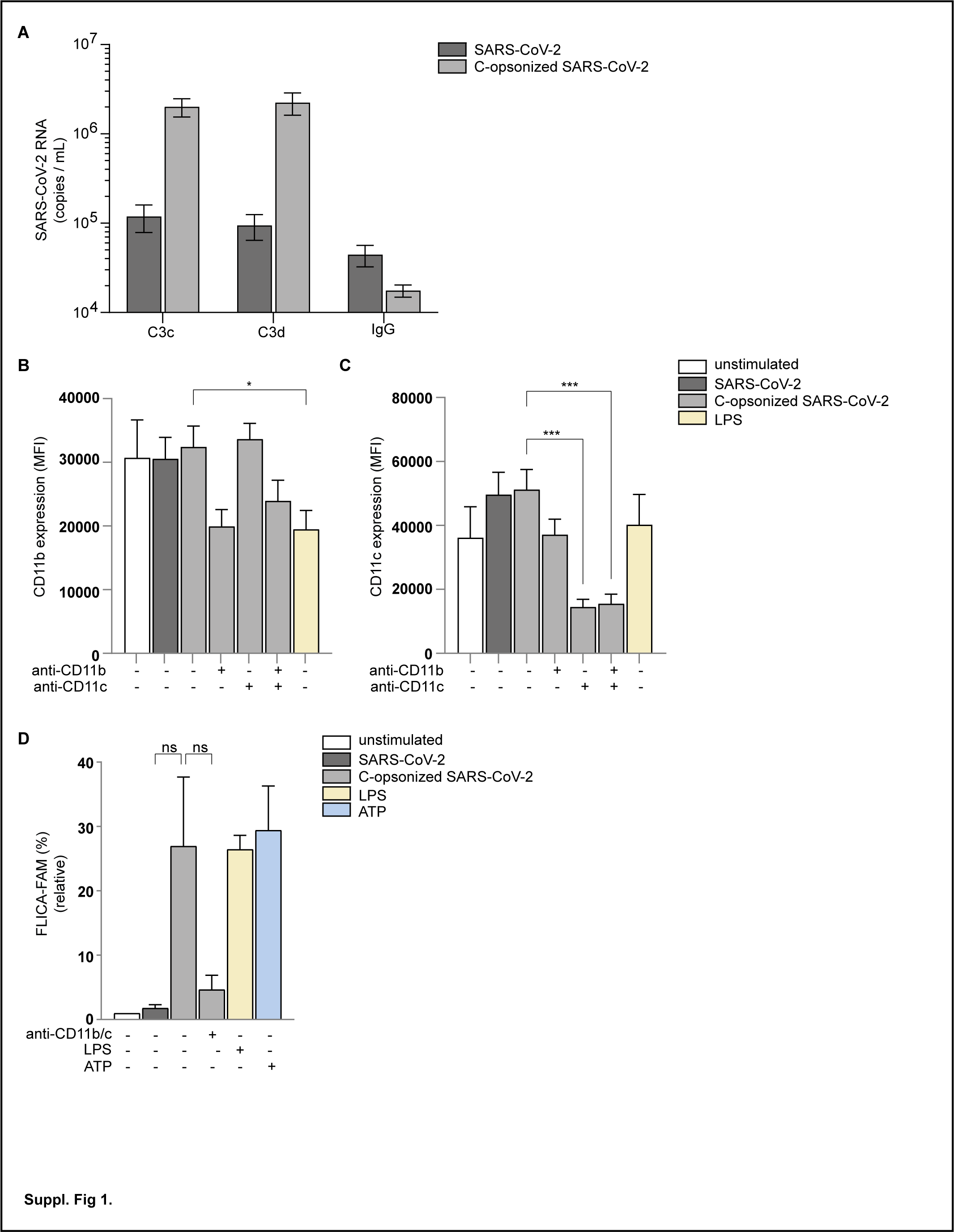

### Suppl. Fig.2

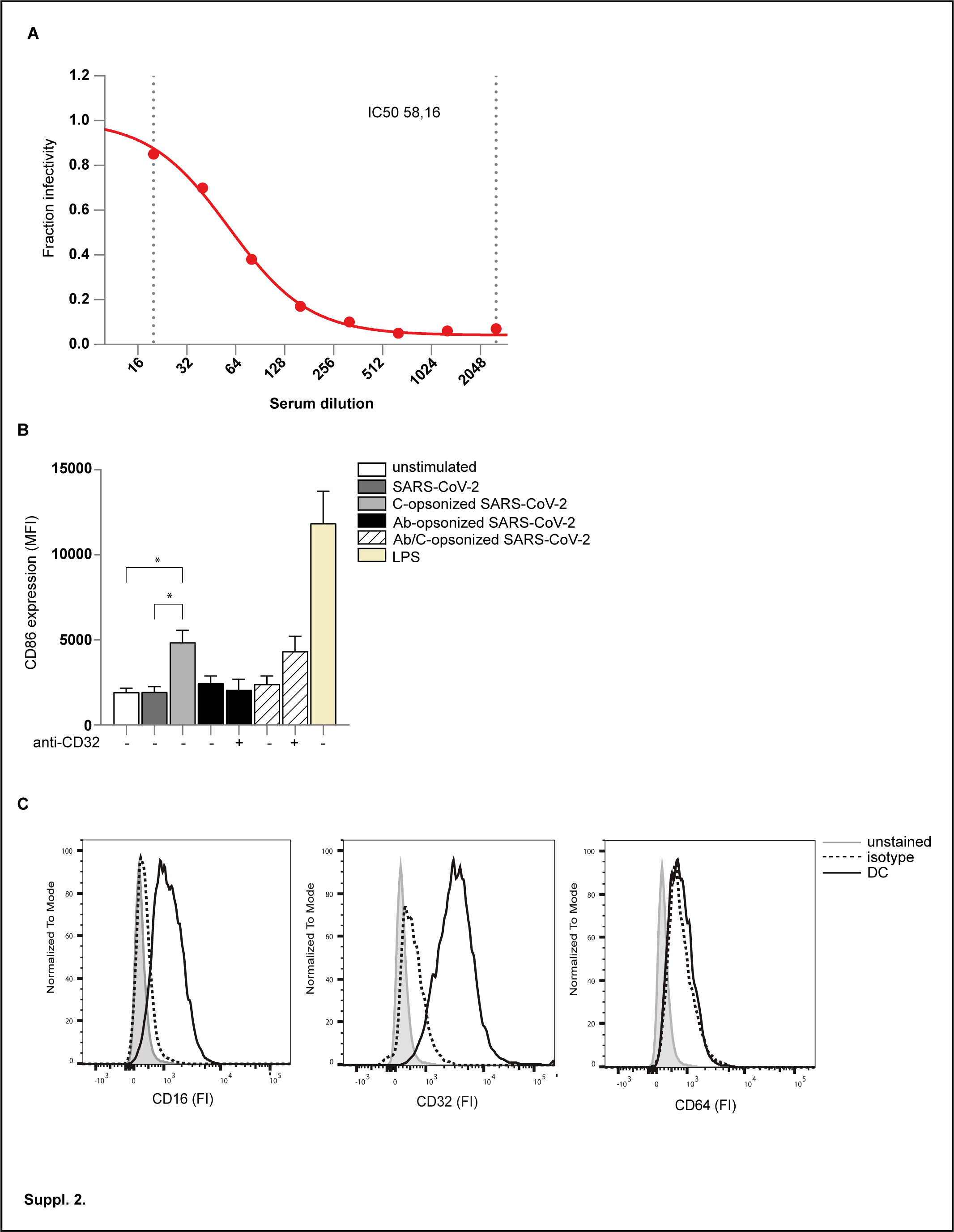
